# Spatial tuning in nociceptive processing is driven by attention

**DOI:** 10.1101/2022.06.16.496352

**Authors:** Wacław M. Adamczyk, Michał Katra, Tibor M. Szikszay, James Peugh, Christopher D. King, Kerstin Luedtke, Robert C. Coghill

## Abstract

When the source of nociception expands across a body area, the experience of pain increases due to the spatial integration of nociceptive information. This well-established effect is called spatial summation of pain (SSp) and has been the subject of multiple investigations. Here, we used cold-induced SSp to investigate the effect of attention on the spatial tuning of nociceptive processing. Forty pain-free volunteers (N=40, 20 females) participated in this experiment. They took part in an SSp paradigm based on three hand immersions into cold water (5°C): Participants either immersed the ulnar segment (“a”), radial segment (“b”) or both hand segments (“a+b”) and provided overall pain ratings. In some trials based on “a+b” immersions, they were also asked to provide divided (i.e., first pain in “a” then in “b”; or reversed) and directed attention ratings (i.e., pain only in “a” or “b”). Results confirmed a clear SSp effect in which reported pain during immersions of “a” or “b” was less intense than pain during immersions of “a+b” (*p*<0.001). Data also confirmed that spatial tuning was altered. SSp was fully abolished when participants provided two ratings in a divided fashion (*p*<0.001). Furthermore, pain was significantly lower when attention was directed only to one segment (“a” OR “b”) during “a+b” immersion (*p*<0.001). We conclude that spatial tuning is dynamically driven by attention as reflected in abolished SSp. Directed attention was sufficient to focus spatial tuning and abolish SSp. Results support the role of cognitive processes such as attention in spatial tuning.

**Perspective:** This article presents experimental investigation of spatial tuning in pain and offers mechanistic insights of contiguous spatial summation of pain in healthy volunteers. Depending on how pain is evaluated in terms of attentional derivative (overall pain, directed, divided attention) the pain is reduced and spatial summation abolished.

## 1. INTRODUCTION

Pain is a complex experience that integrates spatial and temporal aspects of sensory processing. Temporal phenomena such as wind-up have been topics of intensive research. However, the spatial aspects of pain, such as localization^38^, spread^45^, and spatial summation (SSp)^1^, remain poorly understood but have important clinical meaning. For example, the spread of pain is related to disease duration^45^, whereas the size of the painful area contributes to clinical pain intensity^43^.

Computationally, spatial tuning reflects how afferent nociceptive information is integrated within the spatial domain. Spatial tuning can be broad and allow information to be collected from widespread body regions, or spatial tuning could be narrow to optimize the extraction of information from very focal areas. For example, patients with chronic pain have enhanced SSp^16^ and enlarged receptive fields^3^, suggesting a broadening of spatial tuning.

The phenomenon of SSp provides a useful tool for experimental investigation of spatial tuning of nociceptive processing^34^. In the study by Quevedo & Coghill participants provided overall pain ratings, which spatially integrated nociceptive information from two nociceptive foci or directed attention ratings, which were limited to just one of the two simultaneously stimulated foci. Participants were also prompted to divide their attention by first providing a pain rating of one and then the other site. Dividing attention significantly abolished SSp^33,34^. This form of spatial tuning may result from a shrinkage of receptive fields (RF), such that SSp is reduced. This hypothesis is consistent with studies on healthy volunteers^18,34^ as well as awake animals^17^ which showed that nociceptive RFs are shifted or can change their size under the influence of attention. Most importantly, research from other sensory systems supports the role of attention in the spatial shaping of RFs. For instance, a computational modeling study predicts attentionally-driven plasticity of RFs^11^.

Studies of attentionally mediated spatial tuning have only been conducted with noxious heat and in the context of discrete SSp – where spatially separated stimuli are used to evoke pain. Would attentional regulation of spatial tuning also occur in zones of contiguous noxious stimulation? Such regulation may be challenging due to overlapping input from primary afferents. However, experimental demonstration of such regulation would be highly relevant for some types of clinical pain such as complex regional pain syndrome (CRPS), radicular pain or muscle pain such as delayed onset muscle soreness where pain extends in a contiguous fashion^21^. Specifically, disruption of regulation of spatial tuning may contribute to the spread of pain, while restoration of such regulation could provide a novel treatment pathway.

The primary aim of this experiment was to test whether spatial tuning can be attentionally regulated across contiguous zones of noxious stimulation, which has never been addressed. Noxious cold was used instead of heat to investigate if previously observed attentional effects generalize to a different stimulus modality. Lastly, we employed directed-attention ratings to test if attention directed only at a smaller zone of the painful area leads to a systematic reduction in pain. Such conditions have been theoretically associated with a skew of receptive fields towards attended sites^11^ which could also minimize SSp by reducing the efficacy of input from the unattended site.

## 2. METHODS

The study used a within-subject repeated-measurement design. Healthy, pain-free participants underwent multiple Cold Pressor Tasks (CPT) in which they immersed their hand into cold water. The study aimed to assess spatial tuning previously investigated in the context of discrete SSp^34^. Data was collected in a single session (~1.5h duration). The study procedures were approved by the local Bioethical Committee at the Academy of Physical Education in Katowice (1-2021/02-25-2021), and the protocol of the experiment was preregistered in the Open Science Framework using the AsPredicted.org template (https://osf.io/cdwra). The experiment took place in the Laboratory of Pain Research.

### 2.1. Participants

Forty participants with an average age of 27.2 years (± 11.3) took part in the experiment (20 females). Only healthy participants between 19 and 65 years old were included. Participants were excluded if they suffered from chronic (pain lasting >3 months) or acute pain on the study date, took drugs or medications, were diagnosed with a disease related to cold temperature intolerance (e.g., Raynaud syndrome, cryoglobulinemia, cold urticaria, etc.), had experienced in the past a pathological reaction to cold temperature (e.g., excessive edema or redness, blisters, etc.), had a chronic cardiovascular or neurological disease. Recruitment was continued until 40 participants had completed the experiment. Six additional individuals were screened but not included because of not meeting all criteria. All participants completed the entire experiment.

### 2.2. Sample size

This study is considered novel as no previous data on attentionally-driven spatial tuning exists in contiguous SSp with noxious cold as a pain model. The relevant effect size was extracted (Cohen’s *d_z_* = 0.62) from a previous study that used discrete SSp^34^ and reflected the statistical *t* contrast between pain obtained through divided attention ratings vs. overall ratings **(Appendix 1)**. Sample size calculations were performed using the G*Power software^13^ and showed that N=23 was needed to detect the effect with 80% power and that N = 36 was needed to detect the effect with 95% power (α = 0.05_2-tailed_). Because the experimental paradigm presented here differs in terms of the modality as well as the type of pain (CPT, contiguous area of pain) from the study that was used to calculate statistical power^34^, the sample size was increased. Lastly, a sample of 40 prevented us against small group effects that fail to reproduce^6^ and allowed to detect effects as small as *d_z_* = 0.45.

### 2.3. Equipment and CPT

To induce pain, the CPT was employed^25,48^. For this task, participants immersed their dominant hand (or its part) in a circulating cold-water bath UTE-24BB (LaboPlay, Bytom, PL). A fixed temperature of 5°C was maintained by the electronic thermostat of the device as this temperature has been shown to produce moderate pain^25^ in CPT and robust SSp^41^. Pain intensity was rated using the Verbal Rating Scale used previously^42^ which ranges from 0 (no pain at all) to 100 (worst pain imaginable). Participants were instructed that they could provide any number within the 0-100 interval. Skin temperature of the hand was measured before each immersion using a non-contact infrared thermometer (LX-26, ThermoFlash®, Visiomed, FR) to control for confounding effects resulting from variability in skin temperature recovery between immersions. The core of the experiment were whole hand immersions, however, to investigate SSp, only some segments (see **Figure 1**) of the hand were stimulated (see below).

**Figure 1.**
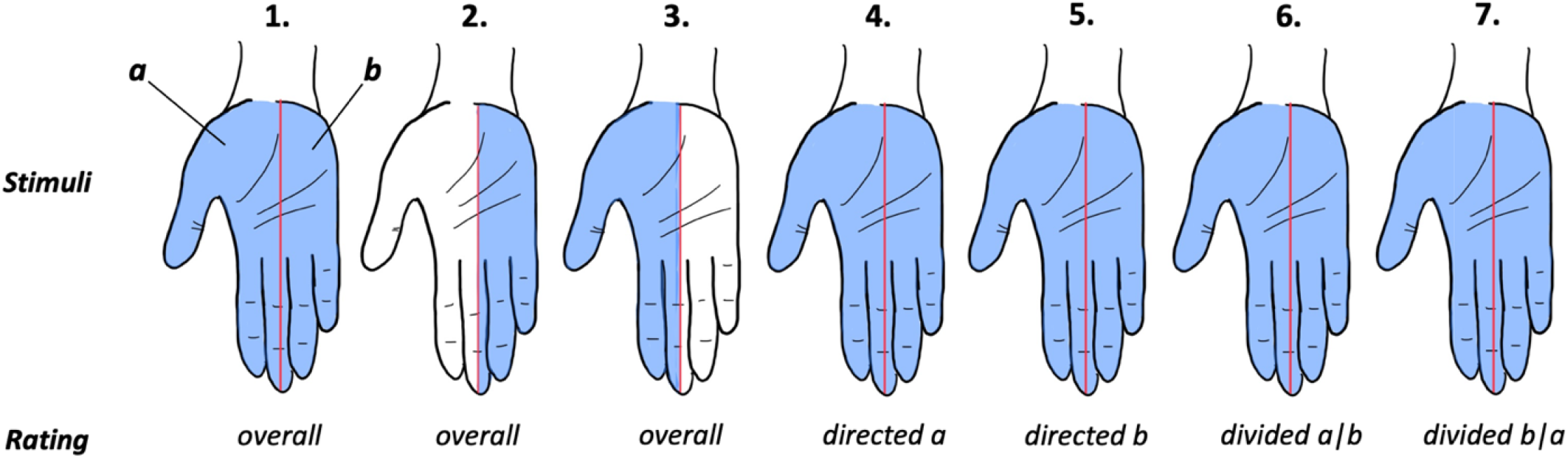
Spatial summation of pain paradigm. Segments of the hand exposed to noxious cold stimulation are presented in blue. The hand was divided into two equal segments (see trial 1), i.e., “a” (radial) and “b” (ulnar). The border dividing the hand into segments was marked on the skin. Participants immersed either two segments simultaneously (“a+b”, trials: 1, 4-7) or just one of them (“a” or “b”, trials 2-3). Overall pain ratings were obtained from both segments (1), only the ulnar (2) or radial (3) segment. Directed attention ratings were obtained from radial (4) and ulnar (5) segments. Divided attention ratings were obtained sequentially such that the first rating from “a” then “b” (trials 6) or in a reversed fashion (trial 7). These trials were presented in two pseudorandom blocks with block I using the trial order of 3-4-1-6-2-7-5 and block II using the trial order 5-1-6-2-4-3-7.

### 2.4. Training phase

Upon arrival at the laboratory and after providing informed consent, participants were screened for a set of inclusion and exclusion criteria. They received an explanation that they would undergo multiple repetitions of 7 different trials (immersions) using CPT. Participants’ hands were divided into two segments: radial (referred to as a segment “a”) and ulnar (referred to as a segment “b”, **Figure 1**). To delineate sections of the hand, an ink-based black line was drawn on the subjects’ hand, starting from the tip of the middle finger up to the distal crease of the wrist. The size of the hand was measured using centimeter tape (length, width, circumference). Following the preparation and anthropometric measures, participants took part in a training session, in which they familiarized themself with the water environment, and most importantly, they were trained on how to perform different types of immersions by pronating and supinating their forearm. Every participant underwent initial training to find a comfortable position for hand placement, and to ensure that half of the hand was stimulated: To do this the forearm was stabilized over the edge of the water bath whereas the top of the middle finger was in a contact with metal tank of the bath. In addition, each brief immersion (30s) was supervised by the examiner and the position of the hand was corrected if necessary. In general, participants’ hand and body position did not deviate from bodily movements occurring in daily life. Apart from immersions, participants were trained in how to provide varied form of pain ratings, i.e., divided attention, directed attention, or overall pain ratings.

During the training session, each type of trial was presented one time (in total 7 trials/immersions), but participants were offered to practice more if needed. Immersions of 30s durations in the training session were separated by 30s breaks to reduce study time and thereby maximize the participants’ attention. Cold pain threshold was measured (i.e., duration of immersion until the pain starts) as a part of the training session and after the main session. After training, a 10-minute break was followed by the experimental session. In the end of the sessions participants completed Pain Vigilance Awareness Questionnaire (PVAQ).

### 2.5. Experimental phase

Following the training session, participants took part in the main session divided into two blocks of 7 trials each. Trials were presented pseudo-randomly within each block. Each trial consisted of a 30s stimulation period (180s interval). Participants fixed their gaze on the middle point of the stimulated area and were prompted to provide pain ratings immediately before hand withdrawal. The following trials were employed **(see Figure 1)**: (1) immersion of two segments, i.e., whole hand (“a+b”) with overall pain rating: Participants were instructed that when prompted (after 25s of immersion) they must provide overall pain from this immersion, (2) immersion of segment “b” and overall rating, (3) immersion of segment “a” and overall rating. For those trials participants were instructed accordingly: *Please provide intensity of experienced pain from 0 to 100.* These trials were utilized to demonstrate SSp.

The trial marked as (4) was based on immersion of both segments (“a+b”) and attention directed at segment “a” and rating only from the segment “a”, whereas trial (5) was an immersion of both segments (“a+b”) and attention directed at segment “b” and rating only from segment “b”. For that trials participants were instructed accordingly: *I will ask you about pain intensity in the segment “a”* [or “b”]. *Focus only on this segment and provide an intensity of experienced pain from 0 to 100.* These trials were utilized to provide a directed attention task.

The trials marked as (6) and (7) were used to investigate if attention towards one segment and then the other one leads to SSp abolishment. In trial (6) immersion of segments “a+b” was performed and divided attention task such that the first rating was provided from segment “a” then from “b”, trial (7) was the same as a trial (6), but first rating from segment “b” was obtained and then from “a”. For those trials participants were instructed: *I will ask you about pain intensity in the segment “a” and then pain in the segment “b”* [or first “b” then “a”]*. Provide intensity of experienced pain from 0 to 100.* These trials were utilized to provide a divided attention task.

The order of trials presentation was organized pseudo-randomly, with different sequence per block: (sequence I: trials 3-4-1-6-2-7-5, sequence II: trial 5-1-6-2-4-3-7). Half of the participants started with sequence I, while the other half began with sequence II.

### 2.5. Single trial design

Each trial started by measuring the skin temperature of the hand. Subsequently, the participant’s position was adjusted according to the type of the trial (in fact only position for trials (2) and (3) required change in the position) and their hand was immersed into water for 30s, with the type of rating(s) provided in the end of the immersion. The hand was then dried using a disposable towel and its palmar side was then placed on the external shield of the water bath (for 60s), which maintained constant warmth sensation. For subsequent 60s the dorsal side of the hand was warmed up with a similar technique, and for the last 60s of the break, participants maintained their hand under the opposite axilla to normalize the temperature of the skin.

### 2.5. Statistical analysis

Descriptive statistics were reported as means or standard deviations (SD). The main analysis plan followed the one reported in the study protocol, i.e., first the repeated measures analysis of variance (ANOVA) was performed to test for the main effect of the factor “trial” (see above trials 1 to 7). If sphericity was violated the Greenhouse-Geisser correction was applied. Two repetitions of each trial were averaged for this analysis. Planned *t-test* comparisons were used to test effects related to SSp, divided attention, and directed attention **(Table 1)**.

**Table 1.** Main contrasts of interests for hypothesis testing.

| Hypothesis | Stimuli (immersion) | Rating | Contrast |
| --- | --- | --- | --- |
| SSp - more intense pain in stimulation if both segments | $a + b$ vs. $a$ | Overall (trial 1) vs. Overall (trial 3) | 1* |
| | $a + b$ vs. $b$ | Overall (trial 1) vs. Overall (trial 2) | 2* |
| | $a$ vs. $b$ | Overall (trial 3) vs. Overall (trial 2) | 3 |
| Divided attention - less intense pain in divided vs. overall ratings | $a + b$ vs. $a + b$ | Overall (trial 1) vs. Divided $a \vert b$ (trial 6) | 4* |
| | $a + b$ vs. $a + b$ | Overall (trial 1) vs. Divided $b \vert a$ (trial 7) | 5* |
| Directed attention - less intense pain in directed vs. overall ratings | $a + b$ vs. $a + b$ | Overall (trial 1) vs. Directed at $a$ (trial 4) | 6* |
| | $a + b$ vs. $a + b$ | Overall (trial 1) vs. Directed at $b$ (trial 5) | 7* |
| | $a + b$ vs. $a + b$ | Directed at $a$ (trial 4) vs. Directed at $b$ (trial 5) | 8 |
| | $a$ vs. $a + b$ | Overall (trial 3) vs. Directed at $a$ (trial 4) | 9 |
| | $b$ vs. $a + b$ | Overall (trial 2) vs. Directed at $b$ (trial 5) | 10 |
| Directed vs. divided attention - exploratory | $a + b$ vs. $a + b$ | Directed at $a$ (trial 4-5) vs. Divided at $b$ (trial 6-7) | 11 |

Letters in column “stimuli” refer to the immersed segments of the hand. Primary comparisons were marked with\*. The remaining (unmarked) comparisons served as additional control or exploratory tests.

Because the skin temperature restorations can vary between individuals, and to rule out that this factor confounded our results, an additional analysis was performed using a Linear Mixed Model (LMM) with “block” (first, second) and “trial” (1 to 7) as within-subject factors and skin temperatures as a covariate. Also, Pearson product coefficients were used to perform exploratory analyses and test if the SSp effect (calculated as a ratio of pain in “a+b” to the mean of pain from smaller areas, i.e., “a” and “b”) was associated with hand size, sensitivity (CPT, pain at baseline) and fear of pain.

To test if baseline sensitivity drives spatial summation, SSp was correlated with mean pain ratings from trials 2 and 3. The reliability of measurement (short test-retest) was calculated for overall pain by correlating data from first and second block using Intraclass Correlation Coefficient (ICC_(3,1)_. All statistical analyses were performed with the IBM Statistical Package for the Social Sciences (SPSS Version 28, Armonk, NY, USA). Visual presentation of the data was performed using GraphPad Prism v.8.0.0 (GraphPad Software, San Diego, California, USA). The original α level was set at *p* < 0.05 and adjusted for multiple testing using Bonferroni correction. Original *p*-values were reported with additional mark of “/” if not significant after correction. Effects sizes higher than 0.14 (*η*^2^_p_), and 0.8 (*d_z_*) were interpreted as large^10^.

## 3. RESULTS

Descriptive statistics for main and secondary variables are presented in **Table 2.** Forty participants (50% females) completed the experiment. As reported in **Table 2,** the average age of the sample was 27.20 years (± 11.30), body mass 71.57 kg (± 13.75), cold pain threshold 20.38 seconds (± 14.23), PVAQ 44.75 (± 7.15), and fear of pain level was 44.55 (± 21.60). Raw data supporting the results are presented in **Appendix 2**.

**Table 2.** Descriptive statistics.

| Variable | Mean | SD |
| --- | --- | --- |
| <u>Participant characteristics</u> |  |  |
| Age [years] | 27.20 | 11.30 |
| Body mass [kg] | 71.57 | 13.75 |
| Height [cm] | 174.05 | 9.16 |
| Hand length [cm] | 19.31 | 1.59 |
| Hand width [cm] | 8.98 | 0.99 |
| Hand circumference [cm] | 20.35 | 2.14 |
| <u>CPT</u> |  |  |
| Before the testing session [sec] | 20.38 | 14.23 |
| After the testing session [sec] | 22.02 | 14.92 |
| <u>Psychological measures</u> |  |  |
| Fear of pain | 44.55 | 21.60 |
| PVAQ | 47.75 | 7.15 |

**Legend:** PVAQ, Pain and Vigilance Awareness Questionnaire, CPT, Cold Pain Threshold, fear of pain was measured on 0 – 100 scale.

### 3.1. Primary analyses

Repeated-measures ANOVA showed a significant effect of the factor “trial”, indicating that the type of trial influenced reported pain (*F*_[4.05,157.91]_ = 8.52,*p* < 0.001, *η*^2^_p_ = 0.18).

#### Spatial summation (Contrasts 1 to 3)

Paired student *t*-tests revealed that the intensity of pain was higher during the immersion of the whole hand (“a+b”) compared to only immersing the radial (“a”, *M*=9.23, 95% [CI: 6.08, 12.37], *t*_[39]_ = 5.93, *p* < 0.001, *d_z_* = 0.93; Contrast 1) or ulnar (“b”, *M*=9.18, 95% [CI: 5.74, 12.62], *t*_[39]_ = 5.40, *p* < 0.001, *d_z_* = 0.85; Contrast 2) segments. No significant difference in pain intensity was found during immersion of the radial (“a”) and ulnar (“b”) segments individually (*M*=-0.05, 95% [CI: −3.21, 3.11], *t*_[39]_ = −0.03, *p* = 0.98, *d_z_* < 0.01; Contrast 3). This set of comparisons confirmed that the SSp effect was able to be reproduced with noxious cold water in support of previous experiments^19,20^ **(see Figure 2 and 3)**. Interestingly, 5 out of 40 participants (12.5%) did not show SSp; instead, they reported lower pain during immersion of the whole hand.

**Figure 2.**
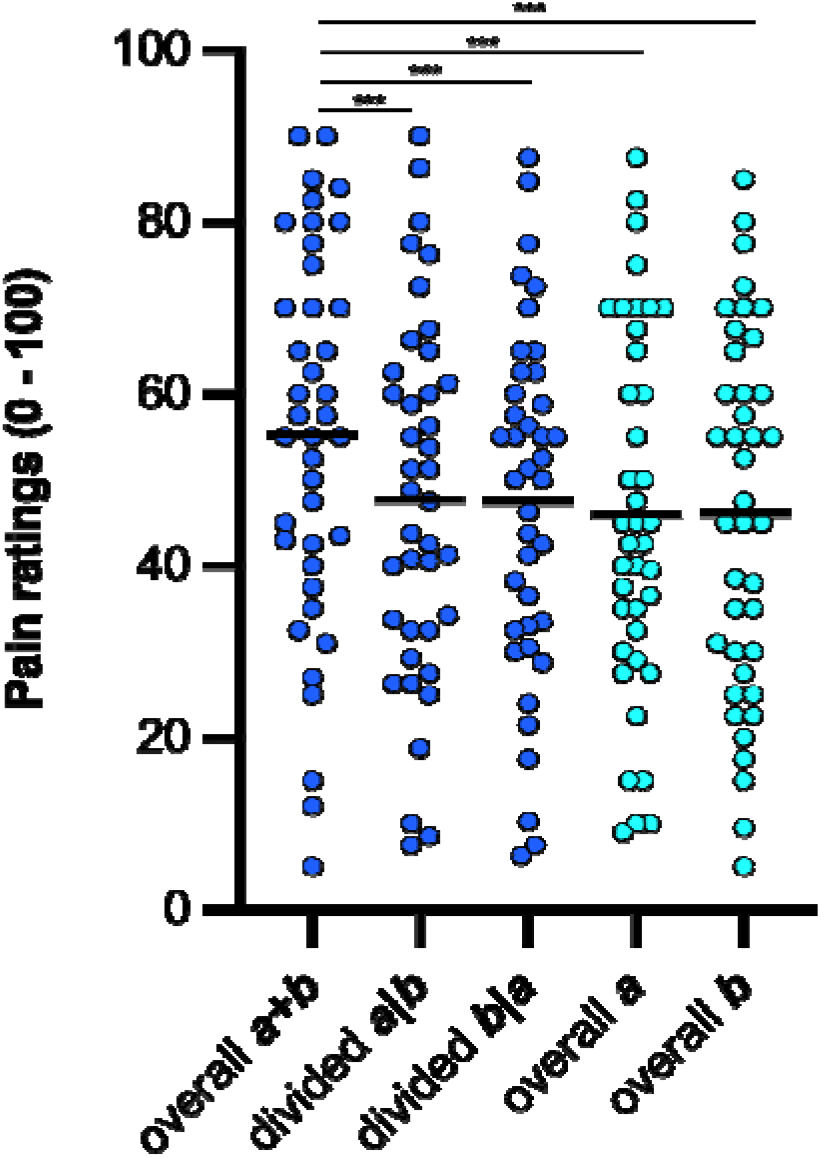
Spatial summation of pain was abolished in the divided attention condition. The horizontal axis includes experimental conditions, and the type of stimuli is marked with colors: Cyan, immersions of segments “a” or “b” alone, dark blue, immersions of segments “a+b”. A clear spatial summation effect was observed when contrasting pain from stimulation of 2 segments (“a+b”) vs. just one segment (“a” or “b”). Note that the spatial summation was fully abolished when ratings were collected in a divided fashion, e.g., first “a” then “b” (“a|b” or reversed). Dots refer to individual data points, horizontal black lines depict mean values. *P*-values are presented following correction: \*\*\**p*< 0.001.

**Figure 3.**
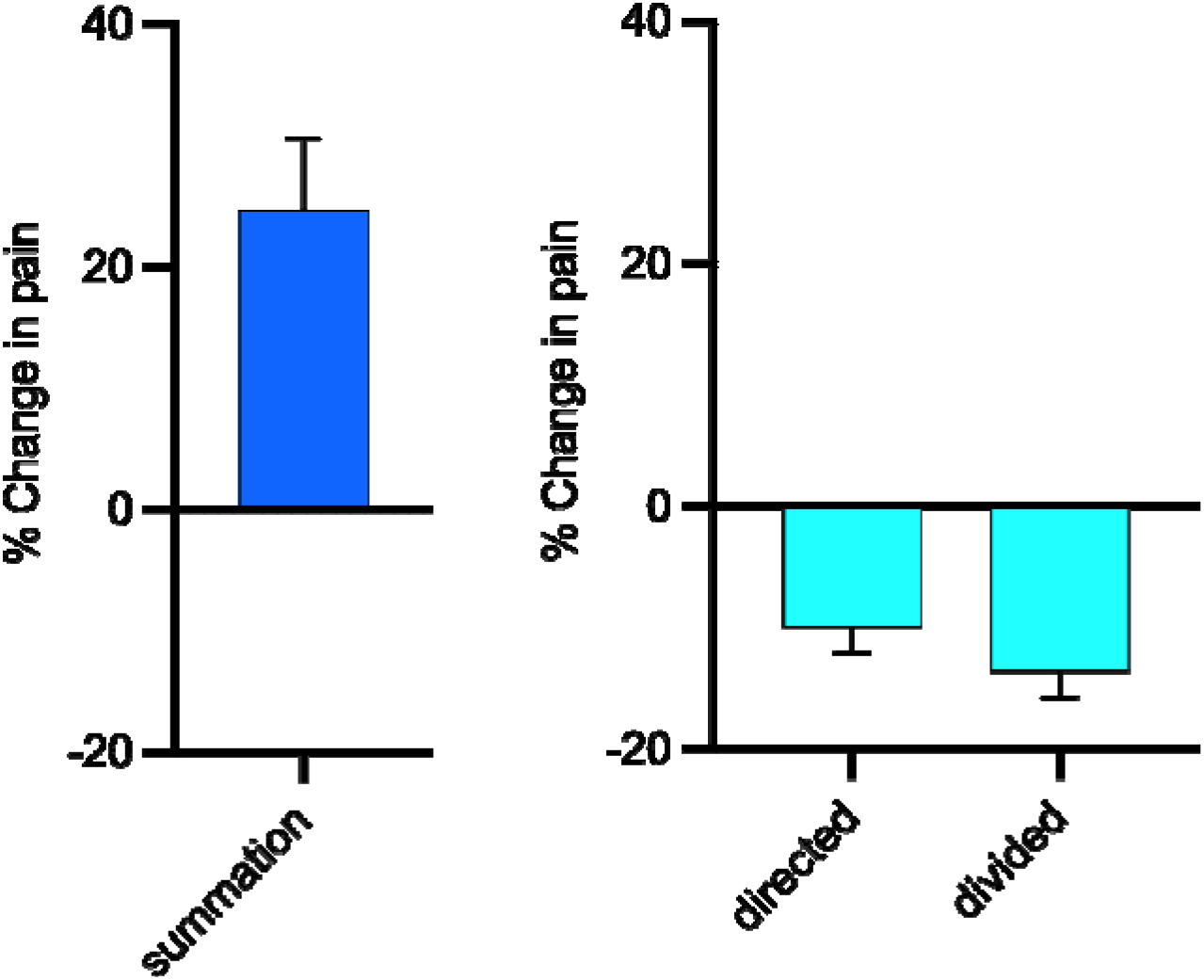
Magnitude of examined effects expressed as relative (%) change in pain. Left panel (summation): Pain increased by 25% during whole hand immersions compared to just one segment. Right panel (attentional effects): Pain decreased by 10% in directed attention ratings and by 14% in divided attention ratings (14%) compared to overall ratings.

#### Divided attention (Contrasts 4 to 5)

Attentional tuning was significant, as SSp was fully abolished when ratings were provided in a divided fashion. Overall pain ratings during the immersion of the whole hand (“a+b”) were significantly higher compared to the same immersion but with divided ratings [i.e., “a|b” (*M*=7.56, 95% [CI: 5.30, 9.81], *t*_[39]_ = 6.78, *p* < 0.001, *d_z_* = 1.07; *Contrast 1)* or “b|a” (*M*=7.74, 95% [CI: 5.15, 10.32], *t*_[39]_ = 6.05, *p* < 0.001, *d_z_* = 0.96; *Contrast 2*)]. Exploratory comparison of directed versus divided attention ratings was not significant (*M* = 1.55, 95% [CI: −0.12, 3.23], *t*_[39]_ = 1.88, *p* = 0.07, *d_z_* = 0.27). These comparisons provide support for attentionally-driven spatial tuning **(Figure 2)**.

#### Directed attention (Contrasts 6 to 8)

*P*ain was perceived as less intense when attention was directed to only one segment (“a” or “b”). When the whole hand (“a+b”) was immersed, pain ratings were less intense in trials with attention directed to only the radial (“a”; *M* = 5.20, 95% [CI: 2.71, 7.69], *t*_[39]_ = 4.23, *p* < 0.001, *d_z_* = 0.67; Contrast 6) or ulnar (“b”;*M*=6.99, 95% [CI: 4.06, 9.92], *t*_[39]_ = 4.82, *p* < 0.001, *d_z_* = 0.76; Contrast 7) segments compared to overall ratings of whole hand immersion (“a+b”). No significant difference in pain was found between attention directed to segment “b” or “a” (*M* = 1.79, 95% [CI: −2.47, 6.04], *t*_[39]_ = 0.85, *p* = 0.40, *d_z_* = 0.13; Contrast 8) during whole hand immersion **(Figure 4)**. Furthermore, directed attention in full hand immersions (“a+b”) led to similar pain level as in immersion involving only segment “b” (*M* = 2.19, 95% [CI: −1.32, 5.70], *t*_[39]_ = 1.26, *p* = 0.22, *d_z_* = 0.20, see **Figure 4**) but not “a” (*M* = 4.03, 95% [CI: 1.06, 6.99], *t*_[39]_ = 2.75, *p* = 0.009/0.049, *d_z_* = 0.43).

**Figure 4.**
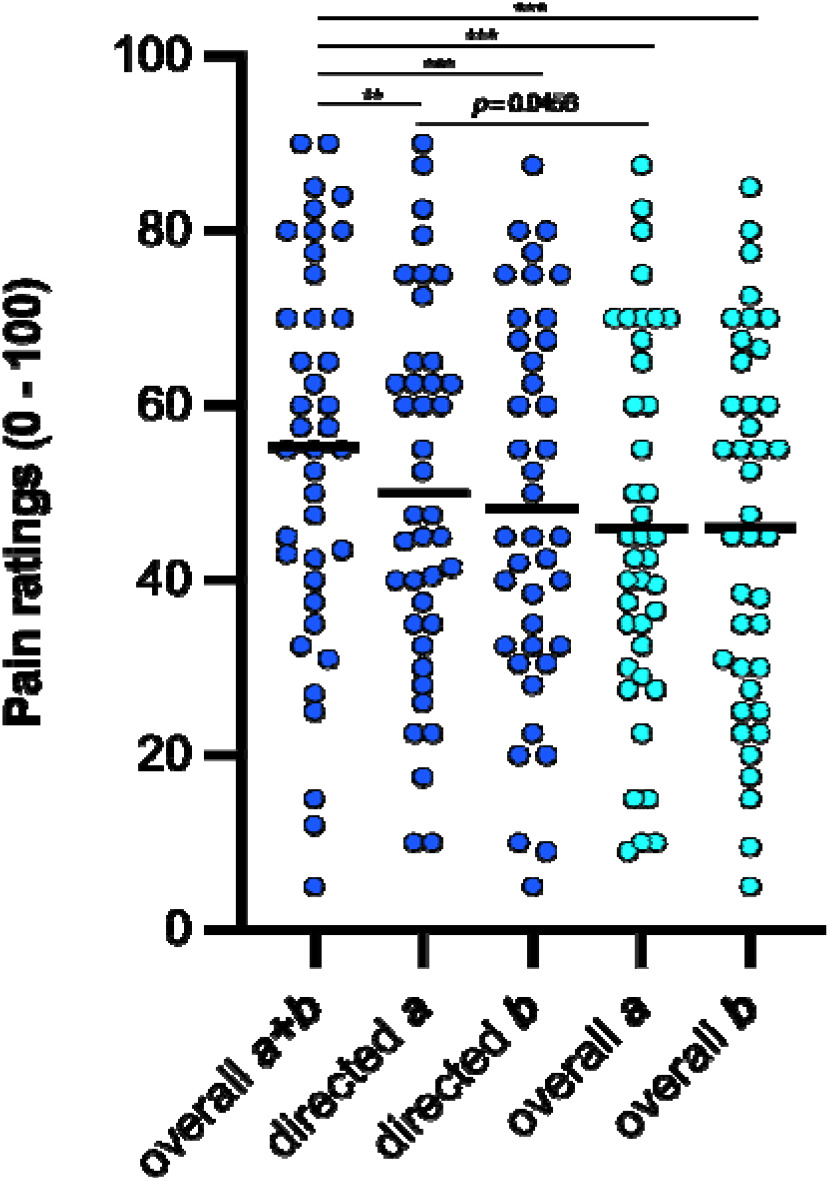
Directed attention diminished experienced pain intensity. The horizontal axis includes experimental conditions, and the type of stimuli is marked with colors: Cyan, immersions of segments “a” or “b” alone, blue, immersions of segments “a+b”. Note that pain was significantly lowered when the two segments were immersed, and participants directed their attention only to one segment of the hand (“a” or “b”). Even though the stimulation area was doubled in these immersions, the pain did not significantly differ from ratings during immersion of only one segment. Dots refer to individual data points, horizontal black lines depict mean values. *P*-values are presented following correction: \*\*\**p*< 0.001, \*\**p*< 0.01.

### 3.2. Additional exploratory analyses

The LMM analysis with skin temperature as a covariate lead to similar results as the original ANOVA. In brief, main effect of “trial”, remained statistically significant (*F*_[6,520]_ = 9.42, *p* < 0.001, *η*^2^_p_), and neither factor “block” (*F*_[1,520]_ = 1.08, *p* = 0.30, *η*^2^_p_ = 0.03) nor “trial” × “block” interaction (*F*_[6,520]_ = 0.43, *p* = 0.86, *η*^2^_p_ = 0.01) was significant. Contrasts corrected by skin temperature are reported in **Appendix 3**. Pain from each trial/immersion was tested for correlation with corresponding measurement of the skin temperature performed right prior to immersion. No significant correlation was found in any of the trials (*r* between 0.15 and 0.28, *p* between 0.08 and 0.48, see **Appendix 4**).

The magnitude of spatial summation (ratio) was not correlated with cold pain threshold measured before (*r* = −0.03, *p* = 0.84) and after the study (*r* = −0.07, *p* = 0.68) and no correlation between SSp and the baseline pain sensitivity (pain in trials 2 and 3) was found (*r* = −0.30, *p* = 0.06). No significant relationship between SSp and fear of pain (*r* = 0.12, *p* = 0.47) or pain hypervigilance (PVAQ, *r* = 0.09, p = 0.57) was found. Anthropometric measurements of the hand such as hand width (*r* = 0.16, *p* = 0.32), length (*r* = 0.11, *p* = 0.51) and circumference (*r* < −0.01, *p* = 0.99) were not related to the magnitude of SSp. Sex differences in the magnitude of SSp (*M*=-0.09, 95% [CI: −0.33, 0.14], *t*_[38]_ = −0.80, *p* = 0.43, *d_z_* = 0.25) and experienced pain ratings (*M* = 2.27, 95% [CI: −12.02, 16.57], *t*_[38]_ = 0.32, *p* = 0.75, *d_z_* = 0.10) were not found. The reliability of pain measurement within the CPT task was good to excellent: ICC_a+b_ = 0.92 (95%CI: 0.85 to 0.95), ICC_a_ = 0.85 (95%CI: 0.73 to 0.92), ICC_b_ = 0.85 (95%CI: 0.74 to 0.92). Note that exploratory analyses are not corrected for multiple testing.

## 4. DISCUSSION

Spatial tuning in the nociceptive system has never been investigated, in contiguous painful area of the body. In the current study attention had an influence on contiguous SSp, which is supported by findings from research on primates ^4^, observations from studies on discrete SSp^33,34^ as well as attentional RF shifts observed in the visual^11^ and auditory systems^14^. The novel observations in our study were that nociceptive foci must not necessarily be spatially distinct for attention to abolish SSp, and that directed attention was sufficient to produce significant pain reductions of the same magnitude as divided attention.

### 4.1. Spatial tuning in acute and chronic pain

At the computational level, spatial tuning reflects how afferent nociceptive information is filtered within the spatial domain. Broad spatial tuning can allow information to be collected from widespread body regions, whereas narrow spatial tuning may extract information from very focal areas. It can constitute the regulation as well as determine the perceptual state. In that sense, the analogy in intensity domain would be adjusting the volume of the sound and then perceiving the chosen loudness. The relatively broad spatial tuning can be responsible for enhanced SSp, whereas focused tuning might effectively hamper SSp such that summation is reduced^16^, disproportionate^22^, nonlinear^1^, or in some circumstances, is even absent^1,36^. At a mechanistic level, tuning may be accomplished by a complex interaction between facilitatory and inhibitory processes that regulate receptive field (RF) sizes of nociceptive neurons^4,9,34^ leading to a determination of spatial attributes of perceived pain.

Previous studies showed that SSp, is amplified in patients with chronic osteoarthritic pain^16^, and fibromyalgia^19^ and that the spatial discrimination of pain may also reflect spatial tuning^26^. In that sense, discrimination may be poorer when spatial tuning is broad and stimulus features cannot be extracted. Although all available studies on chronic pain have been conducted using innocuous stimuli, they provide evidence of disrupted spatial tuning^7^. For example, patients with CRPS^7^, or other types of chronic pain respond poorer with 2-point discrimination tasks^7^. Moreover, more recent studies have confirmed that such a tuning disruption already occurs in the acute phase of pain^27^ or even immediately following noxious stimulus application^2^ and preliminary data shows that intervention directed at spatial retuning may effectively reduce pain intensity^28,29^.

### 4.2. Contiguous SSp

Contiguous SSp induced by cold stimulation was significant and robust (*d_z_*=0.85-0.93), comparable to reports on electrical stimuli^1^, heat^22^, pressure^16^ as well as other studies on noxious cold stimulation^19,20,23,41,47^. Participants immersed one segment (radial/ulnar) or the whole hand into cold water and provided overall pain ratings which were characterized by excellent reliability (ICC ≥ 0.85); pain was greater when a larger area was stimulated. However, the increase in pain was disproportionate: doubling the stimulation area led to a pain increase of only 24%. This is in line with well-documented cases wherein pain produced by the immersion of the entire upper extremity did not double reported pain compared to stimulation of only hand region^22^. Interestingly, the individual differences in the magnitude of SSp showed that in 12.5% of individuals, whole hand immersion produced paradoxically less intense pain, which may be a manifestation of extreme tuning, potentially driven by lateral inhibition^36^.

### 4.3. Attentionally-driven spatial tuning

Pain intensity from two distinct nociceptive foci can be reduced when pain ratings are provided in a divided fashion^34^. In the study by Quevedo & Coghill, divided attention between 2 nociceptive foci (10cm apart) abolished summation, which was explained by the reduction of neuronal output due to RF shrinkage^34^. In addition, attention could disrupt the feeling of connectivity between two stimuli, previously shown to contribute to robust SSp^35^. In a study by Defrin et al.^12^ two discrete heat stimuli were applied wherein participants directed their attention to only one stimulus, and as a result, the overall pain significantly dropped. Thus, directing attention can transform pro-nociceptive into an antinociceptive profile^12^, but more importantly, such a transformation can occur when pain is limited to one body region. Indeed, in this current study, directed attention to just one segment of painful hand significantly abolished SSp.

In some trials of the present study, participants were instructed to divide their attention between different zones of stimulated part of the body. Such a task is likely to be more complex to perform with contiguous stimuli, given the lack of non-stimulated space between nociceptive foci. Nevertheless, despite this difficulty, SSp was abolished in line with previous observations with discrete SSp^34^. An attentional switch between different zones could work as a distraction mechanism leading to pain reduction, however this is not the case as comparable pain reduction was observed in directed-attention conditions where subjects focused on either the ulnar or the radial side of the hand only.

In the previous experiment, directed attention did not produce significant pain reductions^34^, but lower ratings in divided compared to directed attention ratings. In contrast, the current study showed that divided attention has a comparable effect on SSp abolishment as directed attention alone. Yet, the magnitude of the effect was slightly larger in the former (*d_z_*=0.72vs1.02). Thus, directing attention to a small part of the body can inhibit nociception from the surrounding tissue. Indeed, a robust lateral inhibition in the visual system leads to illusionary contrast (e.g. Mach bands) or even abolishment of perceived objects despite their reflection on retina cells^37^. Another relevant visual phenomenon is Troxler fading^44^. Here, presentation of a dot surrounded by shaded low-contrast object fades away after stable gaze fixation at a dot^24^. In the nociceptive system, focusing on the target zone of the painful area might “push” away peripheral zones and thus attenuate pain from surrounding areas.

### 4.4. Attentional regulation of spatial tuning

Our results support the model proposed previously^34^, i.e., total neuronal output was reduced in divided attention ratings because RFs were thought to shrink, in contrast to overall ratings where RFs were thought to enlarge. This is in line with experiments showing that reflex RFs are reduced when attention is directed to a stimulated site of the body compared to a distraction condition, e.g., the Stroop task^18^. The ultimate RF size is likely to be the result of the complex interaction between top-down and bottom-up processes starting from the activation of nociceptive neurons within the dorsal horn. Spinothalamic tract neurons were shown to be involved in a variety of spatial aspects of pain, such as its radiation^32^ and summation^40^. At the supraspinal level ventral and dorsal nociceptive pathways contribute to the perception of different sensory features of pain^30^. The ventral pathway conveys information regarding the intensity of noxious stimuli from the primary (SI) and secondary sensory cortices (SII) to the insula and prefrontal cortex. In contrast, the dorsal subsystem is involved with spatial aspects of pain and conveys information from the somatosensory cortices to the posterior parietal cortex (PPC) and right dorsolateral prefrontal cortex (DLPFC)^30^. Brain imaging studies have shown that the PPC and DLPFC are crucially involved in spatial discrimination of pain^31^. Although SSp with attentional effects has not yet been tested in terms of brain mechanisms, lesion studies confirmed that damage to the PPC can result in excessive pain, spread over large parts of the body^39^, a further manifestation of spatial tuning broadening.

### 4.6. Decomposing pain through attention: clinical implications

One possible non-invasive treatment strategy employed to treat pain is distraction. Distraction involves shifting attention away from pain to another type of stimulation^4^. Although some evidence supports this technique in children^5^, its clinical application to chronic pain populations in adults remains controversial^46^ with potential side effects such as paradoxically higher pain following the distraction procedure^8,15^. Interestingly, it seems that physiologically, reflex RFs become enlarged following a distraction maneuver^18^, whereas they shrink when attention is directed to painful stimulation^4^. Thus, engaging in pain-related attention, a task that can be potentially designed as a mixture of divided and directed attention ratings, may potentially reduce pain, particularly during chronic pain syndromes characterized by extensive pain radiation. Under this framework, the spatial distribution of pain can be decomposed to smaller, less interconnected zones by attentionally mediated engagement of inhibition, leading to pain relief.

### 4.5. Limitations and future directions

Although the paradigm involved novel, feasible techniques of hand immersion, the stimulation itself was not externally controlled like in other studies which utilized stimulator devices (e.g., thermode). Nevertheless, despite some random variation in the stimulation (e.g., bodily and hand position, anthropometric differences), reported pain was greater when a larger area was stimulated, thus reproducing SSp reported previously. Another limitation of the paradigm was its potential for carry-over effects, which -to some extent-has been mitigated by warming hand between trials and controlling for skin temperature. Still, the potential for confounds, such as numbness caused by 5°C stimulation can be reduced even further by employing longer inter-stimulus intervals or reducing the stimulus noxiousness. Furthermore, although two different stimulus sequences were used in the counterbalanced fashion, and no differences in sensitivity were found between them, a fully random order is advised in subsequent studies to rule out potential order effects. Expectations and/or deductive reasoning could have contributed to the reduced pain noted in the divided attention condition. However, in our previous spatial tuning study, explicit controls for expectations confirmed that divided attention itself was sufficient to produce pain reductions. Specifically, manipulating the attentional/cognitive load by adding a neutral probe with 0% probability of activation evoked less inhibition than when the neutral probe had a 50% probability of activation^34^. Thus, whether the pain inhibition in e.g., directed attention condition is solely due to SSp reduction and not because of residual cognitive mechanisms must be determined in subsequent studies. Together with other types of trials, such as ratings of pain while directing attention away from noxious stimulus could also answer the question of how much variability of pain is explained by pure distraction.

## 5. ACKNOWLEDGEMENTS

The study took place in the certified Laboratory of Pain Research (ISO 9001:2015, registration number PW-41212-21K). The authors would like to thank Natalia Kocon for her help with graphical content of the paper.

## 8. APPENDIXES

**Appendix 1.** Extracted data used for sample size calculation.

| Condition | Value extracted | Extracted | SEM | Mean | N | N<sup>(0.5)</sup> | SD |
| --- | --- | --- | --- | --- | --- | --- | --- |
| divided / distal | Mean | 2.733 |  |  |  |  |  |
| divided / distal | SEM | 3.317 | 0.583 | 2.733 | 15 | 3.873 | 2.259 |
| overall / distal | Mean | 4.200 |  |  |  |  |  |
| overall / distal | SEM | 4.850 | 0.650 | 4.200 | 15 | 3.873 | 2.517 |
| divided / proximal | Mean | 3.033 |  |  |  |  |  |
| divided / proximal | SEM | 3.600 | 0.567 | 3.033 | 15 | 3.873 | 2.195 |
| overall / proximal | Mean | 4.217 |  |  |  |  |  |
| overall / proximal | SEM | 4.867 | 0.650 | 4.217 | 15 | 3.873 | 2.517 |
| **Mean overall** | - | - | - | **4.208** | - | - | **2.517** |
| **Mean divided** | - | - | - | **2.883** | - | - | **2.227** |

\*Sample size calculation was made using G\*power for the following comparison: overall pain ratings vs. divided attention pain ratings. Relevant data were extracted from Figure 2 of the study by Quevedo & Coghill \(2007\). Extraction was made using WebPlotDigitizer. Standard errors were converted to SD using the formula. In red means and standard deviations used in sample size calculation.

**Appendix 3.** Contrasts from Linear Mixed Model with temperature as a covariate.

| Trial | CONTRASTS |  |  |  |  |  |  |  |  |  |  |
| --- | --- | --- | --- | --- | --- | --- | --- | --- | --- | --- | --- |
|  | I<sup>***</sup> | II<sup>***</sup> | III<sup>0.97</sup> | IV<sup>***</sup> | V<sup>***</sup> | VI<sup>0.23</sup> | VII<sup>***</sup> | VIII<sup>***</sup> | IX<sup>0.007</sup> | X<sup>0.138</sup> | XI<sup>0.137</sup> |
| 1) a+b overall | **+1** | **+1** | 0 | **+1** | **+1** | 0 | **+1** | **+1** | 0 | 0 | 0 |
| 2) a overall | **-1** | 0 | **+1** | 0 | 0 | 0 | 0 | 0 | **+1** | 0 | **+1** |
| 3) b overall | 0 | **-1** | **-1** | 0 | 0 | 0 | 0 | 0 | 0 | **+1** | **+1** |
| 4) a+b directed a | 0 | 0 | 0 | **-1** | 0 | **+1** | 0 | 0 | **-1** | 0 | **-1** |
| 5) a+b directed b | 0 | 0 | 0 | 0 | **-1** | **-1** | 0 | 0 | 0 | **-1** | **-1** |
| 6) a+b divided a\|b | 0 | 0 | 0 | 0 | 0 | 0 | **-1** | 0 | 0 | 0 | 0 |
| 7) a+b divided b\|a | 0 | 0 | 0 | 0 | 0 | 0 | 0 | **-1** | 0 | 0 | 0 |

**Legend:** Note that the *p* value has been marked in the upper index of each contrast (from I to X). \*\*\* indicate *p* < 0.001. Letters in the first column depict the immersed area of the hand (a, b or a+b). *P*-values were still significant following Bonferroni correction.

**Appendix 4.** Correlations with skin temperatures.

| Trial | *r* coefficient | *p* | *N* | Pain |  | Temperature |  |
| --- | --- | --- | --- | --- | --- | --- | --- |
|  |  |  |  | Mean | SD | Mean | SD |
| 1) a+b overall | 0.17 | 0.28 | 40 | 55.26 | 22.06 | 32.95 | 1.66 |
| 2) a overall | 0.17 | 0.29 | 40 | 46.04 | 21.28 | 32.96 | 1.75 |
| 3) b overall | 0.15 | 0.34 | 40 | 46.09 | 21.08 | 32.39 | 1.62 |
| 4) a+b directed a | 0.28 | 0.08 | 40 | 50.06 | 21.07 | 32.96 | 1.71 |
| 5) a+b directed b | 0.20 | 0.23 | 40 | 48.27 | 21.81 | 32.93 | 1.95 |
| 6) a+b divided a\|b | 0.12 | 0.48 | 40 | 47.71 | 21.10 | 32.50 | 1.91 |
| 7) a+b divided b\|a | 0.13 | 0.43 | 40 | 47.53 | 20.37 | 32.80 | 1.84 |

**Legend:** stimulated segment is marked outside parentheses, whereas rated segment(s) is inside parentheses. Pain was measured using 0-100 Verbal Ratings Scale.

## REFERENCES

1. Adamczyk WM, Manthey L, Domeier C, Szikszay TM, Luedtke K: The nonlinear increase of pain in distance-based and area-based spatial summation. Pain 162:1771–80, 2021.

2. Adamczyk WM, Saulicz O, Saulicz E, Luedtke K: Tactile acuity (dys)function in acute nociceptive low back pain: a double-blind experiment. Pain 159:427–36, 2018.

3. Biurrun Manresa JA, Neziri AY, Curatolo M, Arendt-Nielsen L, Andersen OK: Reflex receptive fields are enlarged in patients with musculoskeletal low back and neck pain. Pain 154:1318–24, 2013.

4. Bjerre L, Andersen AT, Hagelskjær MT, Ge N, Mørch CD, Andersen OK: Dynamic tuning of human withdrawal reflex receptive fields during cognitive attention and distraction tasks. Eur J Pain 15:816–21, 2011.

5. Bukola IM, Paula D: The Effectiveness of Distraction as Procedural Pain Management Technique in Pediatric Oncology Patients: A Meta-analysis and Systematic Review. J Pain Symptom Manage 54:589–600.e1, 2017.

6. Button KS, Ioannidis JPA, Mokrysz C, Nosek BA, Flint J, Robinson ESJ, Munafò MR: Power failure: why small sample size undermines the reliability of neuroscience. Nat Rev Neurosci 14:365–76, 2013.

7. Catley MJ, O’Connell NE, Berryman C, Ayhan FF, Moseley GL: Is tactile acuity altered in people with chronic pain? a systematic review and meta-analysis. J Pain 15:985–1000, 2014.

8. Cioffi D, Holloway J: Delayed costs of suppressed pain. J Pers Soc Psychol 64:274–82, 1993.

9. Coghill RC: The Distributed Nociceptive System: A Framework for Understanding Pain. Trends Neurosci 43:780–94, 2020.

10. Cohen J: Statistical Power Analysis for the Behavioral Sciences. New York: Routledge; 2013.

11. Compte A, Wang X-J: Tuning Curve Shift by Attention Modulation in Cortical Neurons: a Computational Study of its Mechanisms. Cerebral Cortex 16:761–78, 2006.

12. Defrin R, Tsedek I, Lugasi I, Moriles I, Urca G: The interactions between spatial summation and DNIC: effect of the distance between two painful stimuli and attentional factors on pain perception. Pain 151:489–95, 2010.

13. Faul F, Erdfelder E, Lang A-G, Buchner A: G*Power 3: a flexible statistical power analysis program for the social, behavioral, and biomedical sciences. Behav Res Methods 39:175–91, 2007.

14. Fritz J, Shamma S, Elhilali M, Klein D: Rapid task-related plasticity of spectrotemporal receptive fields in primary auditory cortex. Nat Neurosci 6:1216–23, 2003.

15. Goubert L, Crombez G, Eccleston C, Devulder J: Distraction from chronic pain during a pain-inducing activity is associated with greater post-activity pain. Pain 110:220–7, 2004.

16. Graven-Nielsen T, Wodehouse T, Langford RM, Arendt-Nielsen L, Kidd BL: Normalization of widespread hyperesthesia and facilitated spatial summation of deep-tissue pain in knee osteoarthritis patients after knee replacement. Arthritis Rheum 64:2907–16, 2012.

17. Hayes RL, Dubner R, Hoffman DS: Neuronal activity in medullary dorsal horn of awake monkeys trained in a thermal discrimination task. II. Behavioral modulation of responses to thermal and mechanical stimuli. J Neurophysiol 46:428–43, 1981.

18. Henrich MC, Frahm KS, Coghill RC, Andersen OK: Spinal nociception is facilitated during cognitive distraction. Neuroscience 491:134–45, 2022.

19. Julien N, Goffaux P, Arsenault P, Marchand S: Widespread pain in fibromyalgia is related to a deficit of endogenous pain inhibition. Pain 114:295–302, 2005.

20. Julien N, Marchand S: Endogenous pain inhibitory systems activated by spatial summation are opioid-mediated. Neurosci Lett 401:256–60, 2006.

21. Maleki J, LeBel AA, Bennett GJ, Schwartzman RJ: Patterns of spread in complex regional pain syndrome, type I (reflex sympathetic dystrophy). Pain 88:259–66, 2000.

22. Marchand S, Arsenault P: Spatial summation for pain perception: interaction of inhibitory and excitatory mechanisms. Pain 95:201–6, 2002.

23. Martikainen IK, Närhi MV, Pertovaara A: Spatial integration of cold pressor pain sensation in humans. Neuroscience Letters Elsevier Science; 361:140–3, 2004.

24. Martinez-Conde S, Macknik SL, Hubel DH: The role of fixational eye movements in visual perception. Nat Rev Neurosci 5:229–40, 2004.

25. Mitchell LA, MacDonald RAR, Brodie EE: Temperature and the cold pressor test. J Pain 5:233–7, 2004.

26. Mørch CD, Andersen OK, Quevedo AS, Arendt-Nielsen L, Coghill RC: Exteroceptive aspects of nociception: insights from graphesthesia and two-point discrimination. Pain 151:45–52, 2010.

27. Morf R, Pfeiffer F, Hotz-Boendermaker S, Meichtry A, Luomajoki H: Prediction and trend of tactile acuity, pain and disability in acute LBP: a six-month prospective cohort study. BMC Musculoskelet Disord 22:666, 2021.

28. Moseley GL, Wiech K: The effect of tactile discrimination training is enhanced when patients watch the reflected image of their unaffected limb during training. Pain 144:314–9, 2009.

29. Moseley GL, Zalucki NM, Wiech K: Tactile discrimination, but not tactile stimulation alone, reduces chronic limb pain. Pain 137:600–8, 2008.

30. Oshiro Y, Quevedo AS, McHaffie JG, Kraft RA, Coghill RC: Brain mechanisms supporting discrimination of sensory features of pain: a new model. J Neurosci 29:14924–31, 2009.

31. Oshiro Y, Quevedo AS, McHaffie JG, Kraft RA, Coghill RC: Brain mechanisms supporting spatial discrimination of pain. J Neurosci 27:3388–94, 2007.

32. Price DD, Hayes RL, Ruda M, Dubner R: Spatial and temporal transformations of input to spinothalamic tract neurons and their relation to somatic sensations. J Neurophysiol 41:933–47, 1978.

33. Quevedo AS, Coghill RC: An illusion of proximal radiation of pain due to distally directed inhibition. J Pain 8:280–6, 2007.

34. Quevedo AS, Coghill RC: Attentional modulation of spatial integration of pain: evidence for dynamic spatial tuning. J Neurosci 27:11635–40, 2007.

35. Quevedo AS, Coghill RC: Filling-in, spatial summation, and radiation of pain: evidence for a neural population code in the nociceptive system. J Neurophysiol 102:3544–53, 2009.

36. Quevedo AS, Mørch CD, Andersen OK, Coghill RC: Lateral inhibition during nociceptive processing. Pain 158:1046–52, 2017.

37. Ratliff F: Mach Bands: Quantitative Studies on Neural Networks in the Retina. San Francisco: Holden-Day; 1965.

38. Schlereth T, Magerl W, Treede R: Spatial discrimination thresholds for pain and touch in human hairy skin. Pain 92:187–94, 2001.

39. Schmahmann JD, Leifer D: Parietal pseudothalamic pain syndrome: Clinical features and anatomic correlates. Archives of Neurology US: American Medical Association; 49:1032–7, 1992.

40. Sherrington CS: Further observations on the production of reflex stepping by combination of reflex excitation with reflex inhibition. J Physiol (Lond) 47:196–214, 1913.

41. Skalski J, Nastaj J, Swoboda S, Budzisz A, Zbroja E, Malecki, A, Adamczyk, WM: Nielaboratoryjna adap**tac**ja badania przestrzennego sumowania bólu w dobie pandemii COVID-19 [Non-laboratory adaptation to study spatial summation of pain during COVID-19 pandemic]. BÓL 21:1–7, 2021.

42. Staud R, Godfrey MM, Mejia M, Ramanlal R, Riley JL, Robinson ME: Usefulness of Ramp & Hold Procedures for Testing of Pain Facilitation in Human Participants: Comparisons with Temporal Summation of Second Pain. J Pain 21:390–8, 2020.

43. Staud R, Price DD, Robinson ME, Vierck CJ: Body pain area and pain-related negative affect predict clinical pain intensity in patients with fibromyalgia. J Pain 5:338–43, 2004.

44. Troxler, D. (I.P.V.): Über das Verschwinden gegebener Gegenstände innerhalb unseres Gesichtskreises [On the disappearance of given objects from our visual field]. Ophthalmologische Bibliothek 2:1–53, 1804.

45. van Rijn MA, Marinus J, Putter H, Bosselaar SRJ, Moseley GL, van Hilten JJ: Spreading of complex regio**na**l pain syndrome: not a random process. J Neural Transm (Vienna) 118:1301–9, 2011.

46. Van Ryckeghem DM, Van Damme S, Eccleston C, Crombez G: The efficacy of attentional distraction and sensory monitoring in chronic pain patients: A meta-analysis. Clin Psychol Rev 59:16–29, 2018.

47. Westcott TB, Huesz L, Boswell D, Herold P: Several variables of importance in the use of the cold pressor as a noxious stimulus in behavioral research. Percept Mot Skills 44:401–2, 1977.

48. Wolf S, Hardy JD: Studies on pain. Observations on pain due to local cooling and on factors involved in the “cold pressor” effect. J Clin Invest 20:521–33, 1941.

